# oCELLoc: Automated Cell Type Assignment in Transcriptomics Data Using Reference Filtering

**DOI:** 10.64898/2025.12.11.693812

**Authors:** Afeefa Zainab, Vladyslav Honcharuk, Alexis Vandenbon

**Affiliations:** Laboratory of Tissue Homeostasis, Institute for Life and Medical Sciences, Kyoto University, Kyoto, 606-8507, Japan; Department of Neuroscience, Graduate School of Medicine, Kyoto University, Kyoto, 606-8507, Japan; Institute for Liberal Arts and Sciences, Kyoto University, Kyoto, 606-8507, Japan; Graduate School of Biostudies, Kyoto University, Kyoto, 606-8507, Japan

## Abstract

Interpreting single-cell RNA sequencing (scRNA-seq) and spatial transcriptomics (ST) data requires accurate cell-type prediction, which strongly depends on the quality of the reference used. However, prediction accuracy is highly dependent on reference quality; missing relevant cell types or including irrelevant ones can substantially impair performance. To address this challenge, we developed oCELLoc, a regression-based method that selects the most appropriate reference cell types from a large atlas and tailors them to each new sample. oCELLoc takes pseudobulk gene expression from ST or scRNA-seq data together with a broad reference matrix and uses regularized regression with cross-validation to identify a limited number of essential cell types. We applied oCELLoc to toy datasets, scRNA-seq data, and 2,144 Visium samples across diverse tissues and conditions, demonstrating that using the filtered cell types leads to more biologically meaningful downstream predictions. oCELLoc is available as an R package on GitHub and CRAN.

## Introduction

Recent developments in ST and scRNA-seq have made it possible to profile gene expression at the cellular level while maintaining spatial context, which has significantly improved our comprehension of tissue organization in both health and illness. While scRNA-seq captures transcriptomes at individual cell resolution, it also detects cell types and dynamic cell states (1), ST maintains the positional context of expression across tissue sections, mapping the spatial distribution of cells and their expression patterns (2). Our understanding of tissue architecture and function in both health and illness has been completely transformed by these beneficial techniques working together. Assigning expression to certain cell types is made more difficult by the fact that ST techniques frequently measure aggregate RNA from spots that may contain many cells (with resolution varying by platform, in some cases from near single-cell up to tens or hundreds of cells per spot). Separating signals from mixtures of different cell types is necessary for cell type prediction or deconvolution in transcriptomic data, whether from scRNA profiles or ST data. Using single-cell reference datasets that capture distinctive cell-type signals of gene expression, a variety of computational approaches have been developed for evaluating cell-type composition, typically by using the cell-type-specific transcriptome profiles obtained from an appropriate single-cell reference to deconvolve cell-type mixtures (3-5). Some of these methods for ST include Robust Cell Type Decomposition (RCTD) (6), cell2location (7), SPOTlight (4), spatialDWLS (8), STdeconvolve (9), and various other methods (10) that deconvolves ST data for cell type prediction. Methods used for scRNA data include SingleR (11), BayesPrism(12) and DWLS (13). Generally, reference-based deconvolution techniques require well-annotated reference data to predict accurate cell types in both ST and scRNA deconvolution.

A wide range of techniques based on regression, probabilistic modelling, Bayesian frameworks, nonnegative matrix factorization, and deep learning are being used in the quickly growing field of spatial transcriptomics deconvolution. The variety of approaches and difficulties in the field (14) are highlighted by a recent review of cell-type deconvolution in spatial transcriptomics (10). One general limitation is that spatial deconvolution is generally an ill-posed problem (i.e. numerous alternative solutions) due to significant collinearity among cell type profiles, measurement noise, platform-specific variances, and limited gene coverage per spot. To address this, compressed sensing concepts or sparse regression (e.g. via Lasso) have been proposed, which enforce parsimony (i.e., choosing only a few contributing cell types) to stabilize inference (15). Some published techniques already make use of shrinkage-based or Lasso-based selection. For instance, Lasso is used by SPADE (16) to determine the sorts of cells that are present in a domain (prior to deconvolution). While some approaches explicitly simulate platform effects or use spatial priors (e.g., graph networks), they frequently still assume a fixed reference and do not try an upstream cell-type pruning step. For example, STdGCN (17) integrates spatial neighbors’ information with expression profiles using graph convolutional networks.

Although these promising techniques are proposed for cell type prediction and deconvolution using a reference dataset, an overlooked problem is the selection or construction of the reference dataset itself. The quality of predictions naturally depends on the quality and accessibility of cell type annotations and scRNA-seq data. If relevant cell types (i.e., those present in the target sample we want to predict) are missing from the references, this leads to incorrect predictions (14, 18). Additionally, the inclusion of irrelevant cell types in the reference is likely to negatively impact downstream predictions. In many cases, it is not obvious which cell types are present in a sample, and even for an expert in the field, it might not be easy to classify which cell types should be included in a reference and which should not. Therefore, there is an urgent need for a suitable recommendation on how to optimize the usefulness of current datasets to create reference datasets, given the increasing quantity of scRNA-seq datasets being generated (18).

To overcome these difficulties, we present oCELLoc, a computational tool that aims to carefully determine the cell types found in a transcriptomic dataset. This allows for the creation of a refined reference with an optimized cell type composition according to the given dataset, thus improving downstream cell type deconvolution. In brief, oCELLoc uses shrinkage methods (including Lasso regression, Ridge regression, and Elastic Net) to predict cell types most likely to be present in each target dataset. To do so, oCELLoc takes as input the target dataset (after conversion to pseudobulk data) and a large reference atlas of gene expression profiles of a wide variety of cell types. Shrinkage methods are used to decompose the pseudobulk target data into a small number of cell types. Cross-validation is used to estimate a suitable cutoff threshold for optimal cell types. The advantages of oCELLoc are that it 1) selects relevant cell types/states, 2) excludes irrelevant cell types/states, and 3) limits the computational cost of the downstream deconvolution methods by making this pre-selection (i.e., it’s less time-consuming to run cell type prediction/deconvolution using a reference consisting of fewer cell types). We illustrate the use of oCELLoc by applications on synthetic data with known cell compositions, as well as single-cell and a large collection of spatial transcriptomics data covering a wide variety of tissues and conditions.

## Methods

### oCELLoc methodology

oCELLoc is a computational tool that is designed to facilitate the selection of appropriate reference cell types for deconvolution and cell type prediction in spatial and single-cell transcriptomics data (Figure 1). It requires two main datasets as input: (i) a target dataset (scRNA or ST) in the form of pseudo-bulk expression, representing the combined transcriptional profile of cells or spatial spots respectively for which the significant cell type composition is unknown, and (ii) a comprehensive reference dataset that should contain the average expression profiles of a large and diverse collection of well-annotated cell types across various tissues. oCELLoc uses a shrinkage method that creates sparsity by penalizing the inclusion of irrelevant variables to determine which subset of reference cell types best explains the target dataset. All models are fit using the *glmnet* regularized regression framework (19), which uses a mixing parameter α to control the type of penalty applied: α = 1 for Lasso (L1) (20), α = 0 for Ridge (L2) (21), and intermediate values (0 < α < 1) for Elastic Net (22) (a combination of L1 and L2). As a result, the model automatically narrows the vast reference set to a smaller list of potential cell types that best match the target expression profile.

**Figure 1:**
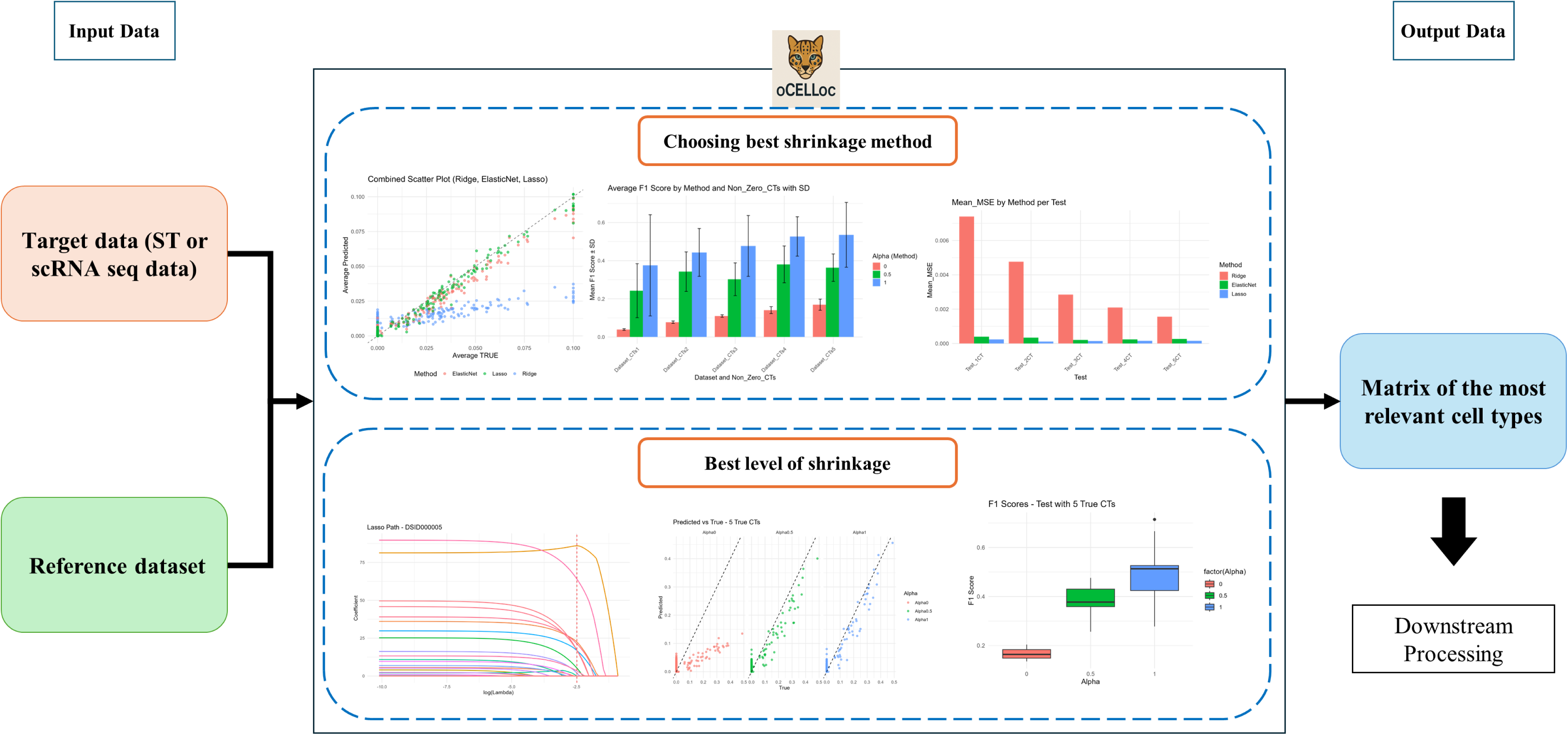
Overview of the oCELLoc workflow. Using test data (ST or scRNA-seq) and a reference matrix as input, oCELLoc assesses various shrinkage regression techniques (Lasso, Ridge, and Elastic Net) and determines the optimal technique and shrinkage level (λ) to get the matrix of the most relevant cell types as an output. Only the most essential cell types are included in the filtered matrix that is produced for further downstream analysis.

#### Model Formulation

In oCELLoc, we averaged the expression across the target dataset S (G genes) to obtain a single large pseudobulk. We also prepared a reference dataset ***R*** with gene expression profiles for various cell types (G genes × C cell types). Each column of ***R*** represents a specific cell type’s average gene expression profile. Only the genes that ***S*** and ***R*** have in common are retained to ensure compatibility. To calculate the contribution of each reference cell type to the observed pseudobulk expression, oCELLoc uses regularized regression for every target sample. The general optimization problem that is resolved is:

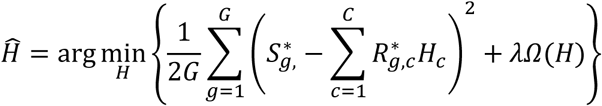

where *S_g_* is the expression of gene *g* in a sample, *R_g,c_* is the expression of gene g in cell type c, *H_c_* is the estimated contribution of cell type c to the target sample, λ is a regularization parameter promoting sparsity. Only non-negative and non-zero values are kept in the prediction matrix. The estimated contributions 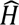 are normalized such that 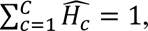 yielding the final cell type proportion matrix 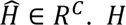 is the vector that has the contribution of cell types to the sample. In conclusion, *H* is the sample-level vector of cell-type contributions that best reconstructs the target pseudobulk expression as a weighted combination of the reference profiles. The penalty term Ω(*H*) determines the regression type:

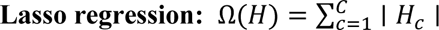

It selects the most relevant cell types by keeping irrelevant cell types close to or at zero.

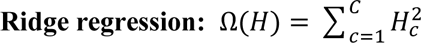

It achieves stability and uniformity by reducing all of the coefficients to zero without imposing sparsity.

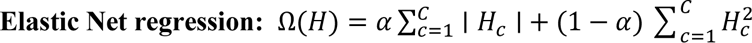

Combining the benefits of both Lasso and Ridge, where 0 ≤ *α* ≤ 1 regulates the ratio of shrinkage to sparsity.

We used cross-validation via cv.glmnet (19) to find the optimal regularization parameter λ* for each ST sample. λ* is chosen as the value of λ that minimizes the mean cross-validation error. The Lasso model is re-fitted using λ* to determine the final cell types present in each sample. We also impose restrictions on the minimum and maximum number of non-zero cell types that can be included in the model to avoid overfitting or under-selection. As output, oCELLoc produces (i) a ranked list of chosen cell types, (ii) the regression coefficients that correspond to those cell types, (iii) the optimal value of the regularization parameter, and (iv) other diagnostic metrics that provide further insight into the robustness and predictive power of the model.

Precision, recall, F1 score, mean absolute error (MAE), and mean squared error (MSE) (23) were used to evaluate performance after predicted coefficients were compared to known cell type proportions. The F1 score is an indicator that goes from 0 to 1, with higher scores signifying a better overall balance between recall and precision. Furthermore, for each dataset and parameter combination, cross-validation curves, coefficient histograms, and metric summaries were produced, making it possible to determine the most suitable regularization parameters for accurate cell type prediction. The metrics listed below were calculated:

▪ Precision = TP / (TP + FP)
▪ Recall = TP / (TP + FN)
▪ F1 = 2 × (Precision × Recall) / (Precision + Recall)
▪ MAE = (1/n) Σ |yᵢ – ŷᵢ|
▪ MSE = (1/n) Σ (yᵢ – ŷᵢ)²

Where TP represents true positives (i.e., a cell type correctly predicted to be present), FP represents false positives (i.e., a cell type incorrectly predicted to be present), and FN stands for false negatives (i.e., a cell type mistakenly predicted to be absent). y_i_ and ŷᵢ denote the actual and predicted proportions of cell type i, respectively. Higher F1 scores, which range from 0 to 1, indicate a better balance between recall and precision. MAE and MSE are used to measure the difference between the true and predicted cell type proportions. MAE calculates the average magnitude of absolute errors between predicted (ŷᵢ) and actual (yi) values, providing an overall assessment of prediction accuracy. A lower MAE suggests that the predicted proportions are closer to the true values on average. MSE, on the other hand, computes the average of the squared differences between predicted and actual values, giving more weight to larger errors. This makes MSE more sensitive to outliers and significant deviations. Overall, these metrics offer a comprehensive evaluation of oCELLoc’s ability to accurately predict underlying cell type composition.

Based on the output of oCELL, a refined reference is created using only the cell types predicted to be present according to oCELLoc. This revised reference is subsequently used for cell type prediction techniques like RCTD (6), SingleR (11), and other deconvolution frameworks. In this way, oCELLoc acts as a pre-processing stage that enhances the precision and comprehensibility of ensuing analysis. To make the methodology available to the community, an R package version of oCELLoc has been created (https://github.com/afeefa-zainab/oCELLoc).

### Preparation of cell type references

We prepared human and mouse references as follows. We collected large reference scRNA datasets from the Tabula Sapiens study (24) that included cell types from 75 tissues from 28 different organs from CellxGENE (25, 26). To these, we further added several missing cell types, including cell types for cancer (tumor cells, abnormal cells, and neoplastic cells) as well as several other tissue-specific cell types, including malignant cells from breast cancer tissue (27), neoplastic cells from liver cancer tissue (28), osteoclasts and osteoblasts from human quadriceps tendons (29) and brain and neuronal cells (30) from the Cellxgene database. The final human reference dataset included 170 cell types. In this dataset, we calculated highly variable genes using Seurat (31) Subsequently, we extracted the average expression data for the 2500 most variable genes for all 170 cell types, thus creating a large pseudobulk matrix (genes x cell types). Similarly, we prepared a reference for 125 mouse cell types. We used the Tabula Muris study (32), which covers 23 different tissues and organs.

### Benchmarking with toy datasets

We first used oCELLoc on toy datasets with known cell type proportions to assess our approach’s performance under controlled conditions.

#### Data Preparation

We created artificial pseudobulk expression datasets as follows. Artificial references were generated containing artificial cell types and their gene expression values for 50 artificial genes. Artificial pseudobulk datasets were generated using a predetermined number of cell types (k = 1 to 5) chosen from the reference datasets described above. The proportions of cell types were generated using random numbers from a uniform distribution (between 0 and 1) and normalized to sum to 1. The appropriate rows of the reference matrix were linearly combined with these proportions to create a synthetic test sample with a known ground-truth composition. To this, noise was added using a normal distribution N(μ=0, *σ*=0.05). For each number of cell types k, 10 different benchmark datasets were generated.

#### Application of oCELLoc

oCELLoc was applied to each toy dataset using the three regression models (Lasso, Ridge, and Elastic Net) to estimate the present cell types. To evaluate the model’s stability and robustness, cross-validation curves, coefficient histograms, scatter plots of expected and true cell types, and metric summaries were generated for each dataset and parameter value. Cross-validation was used to estimate the ideal regularization strength (λ), and the model identified the primary contributing cell types and their coefficients.

#### Accuracy evaluation

The projected coefficients were compared to the actual known proportions that were utilized to create toy datasets to measure the prediction accuracy. Accuracy was measured using metrics described above.

### Applications to single-cell RNA-seq datasets

We used a well-studied single-cell RNA-seq dataset (PBMC3k) (33) to test the applicability of our method on single-cell transcriptomics data.

#### Data preparation

First, standard preprocessing procedures using Seurat were applied to normalize and conduct quality-control, which removal of genes expressed in fewer than 3 cells and cells with fewer than 200 expressed genes. We further normalized data, analyzed highly variable genes using FindVariableFeatures, and scaled data using ScaleData from Seurat (version 5.1.0) (31). We processed this dataset into a pseudobulk dataset by calculating the average expression for each gene across all cells and used this as input to oCELLoc. As a reference, we used the human reference dataset described above as an input to singleR.

#### Application of oCELLoc

Lasso regression, the best-performing shrinkage technique in the toy datasets benchmarks, was used to analyze the pseudobulk expression data with oCELLoc.

#### Accuracy evaluation and downstream annotation

The most suitable cell types chosen by oCELLoc were given as input to SingleR for the deconvolution process. The accuracy of predictions was evaluated by comparing predicted cell types with known annotations in the PBMC3k dataset, using the accuracy metrics described above. To verify robustness, cross-validation curves, coefficient histograms, and metric summaries were generated for every parameter value, just like in the benchmarking of the toy datasets.

### Applications for spatial transcriptomics datasets

#### Data preparation and oCELLoc application

We obtained 10X Genomics Visium data (human: 1361 samples; mouse: 783 samples) from DeepSpaceDB (34, 35), including a wide variety of tissues. A list of sample IDs and their annotations is available in the supplementary Table S1. Each Visium sample was processed into a pseudobulk sample by averaging the expression over all spots. Subsequently, we predicted the present cell types in each sample using oCELLoc.

#### Accuracy evaluation and downstream annotation

oCELLoc was applied to each ST sample with different shrinkage values, using cross-validation to select the optimal shrinkage value. Subsequently, oCELLoc was applied with this optimal shrinkage value to predict cell types in each sample.

## Results

In this study, we introduce oCELLoc, a method that predicts the cell types that are present in a transcriptomics dataset, to aid the downstream cell type prediction process. A typical use case is shown in schematic Figure 1. Users have a single-cell or spatial transcriptomics dataset in which they want to predict cell types. This is typically done based on a reference dataset that includes gene expression profiles of known cell types. However, it is often not clear which cell types are present and absent in the sample, making the choice of a good reference difficult. Any reference might include irrelevant cell types and/or lack necessary cell types, resulting in incorrect predictions. To address this issue, users can use oCELLoc to first select the subset of cell types that appear to be present in the input sample. In brief, oCELLoc applies regularization techniques to select only present cell types and exclude irrelevant cell types. This selection step improves the quality of the downstream cell type prediction or deconvolution.

### Application on toy datasets

To explore the behavior of oCELLoc, we first applied it to toy datasets with known cell type proportions. These toy datasets were created by merging 1 to 5 known cell-type expression profiles with predetermined proportions to create realistic toy pseudobulk samples. Subsequently, oCELLoc was applied on these toy datasets using three different regression models (Lasso, Ridge, and Elastic Net) and using two different λ selection strategies: (1) “Fixed λ ”, which applied a uniform sequence of increasing regularization strengths (for example, from 0 to 1 with 10–20 intervals), and (2) “Auto λ ”, which used cross-validation to automatically select the best λ freely for each method (using cv.glmnet) without giving it any range. This was repeated ten times, and average tendencies were inspected.

The identification of the true contributing cell types from synthetic pseudobulk expression data was consistently better with Lasso (L1) than with Ridge (L2) and Elastic Net (L1+L2) in our fixed λ experiments (Figure 2). As can be seen from the comparison boxplot of F1 scores (Figure 2a and b), Lasso performed best where regularization that induces sparsity is most helpful, achieving the highest F1 Scores over a range of sparsity levels (1–5 true cell types), giving fewer false positives and mostly predicting true cell types. The results of all accuracy matrices are shown in Supplementary Table S2. This dominance is graphically demonstrated in the corresponding scatter plots (Figure 2b), where Ridge and Elastic Net provide lower scores and retain more irrelevant cell types. From Figure 2b, we can see that Lasso and Elastic Nets in general show similar tendencies, with returned coefficients being correlated with the true composition of the toy datasets. In contrast, coefficients predicted by Ridge regression were less correlated with the true compositions. Moreover, Ridge regression and, to a lesser extent, Elastic Nets predicted false positives, i.e., they often predicted non-zero coefficients for absent cell types (true proportion 0). This was less the case for Lasso regression. This behavior results from the way these techniques penalize coefficients: Ridge employs an L2 penalty, which reduces coefficients in the direction of zero but seldom makes them exactly zero. Elastic Net permits modest non-zero coefficients since it mixes L1 and L2 penalties. Lasso regression, on the other hand, uses a pure L1 penalty that encourages sparsity and can drive irrelevant coefficients precisely to zero. Lasso thus generated significantly fewer false positives In the auto-λ tests, where λ for each shrinkage method was chosen automatically using cross-validation, Lasso once more demonstrated the best performance (Figure 2c and d), this time with a better balance between precision and recall, which increased the total F1 Scores shown in boxplots and scatterplots. Even when λ is adjusted separately for each dataset, Lasso’s capacity to reduce noise and concentrate on the right contributors is supported by the scatter plot (Figure 2d). Under auto-λ conditions, Elastic Net performance was slightly improved. On the other hand, Ridge regression performance was consistently bad in both MAE and F1 Score due to many false positives (i.e., non-zero coefficients predicted for irrelevant cell types); the results are shown in Supplementary Table S3.

**Figure 2:**
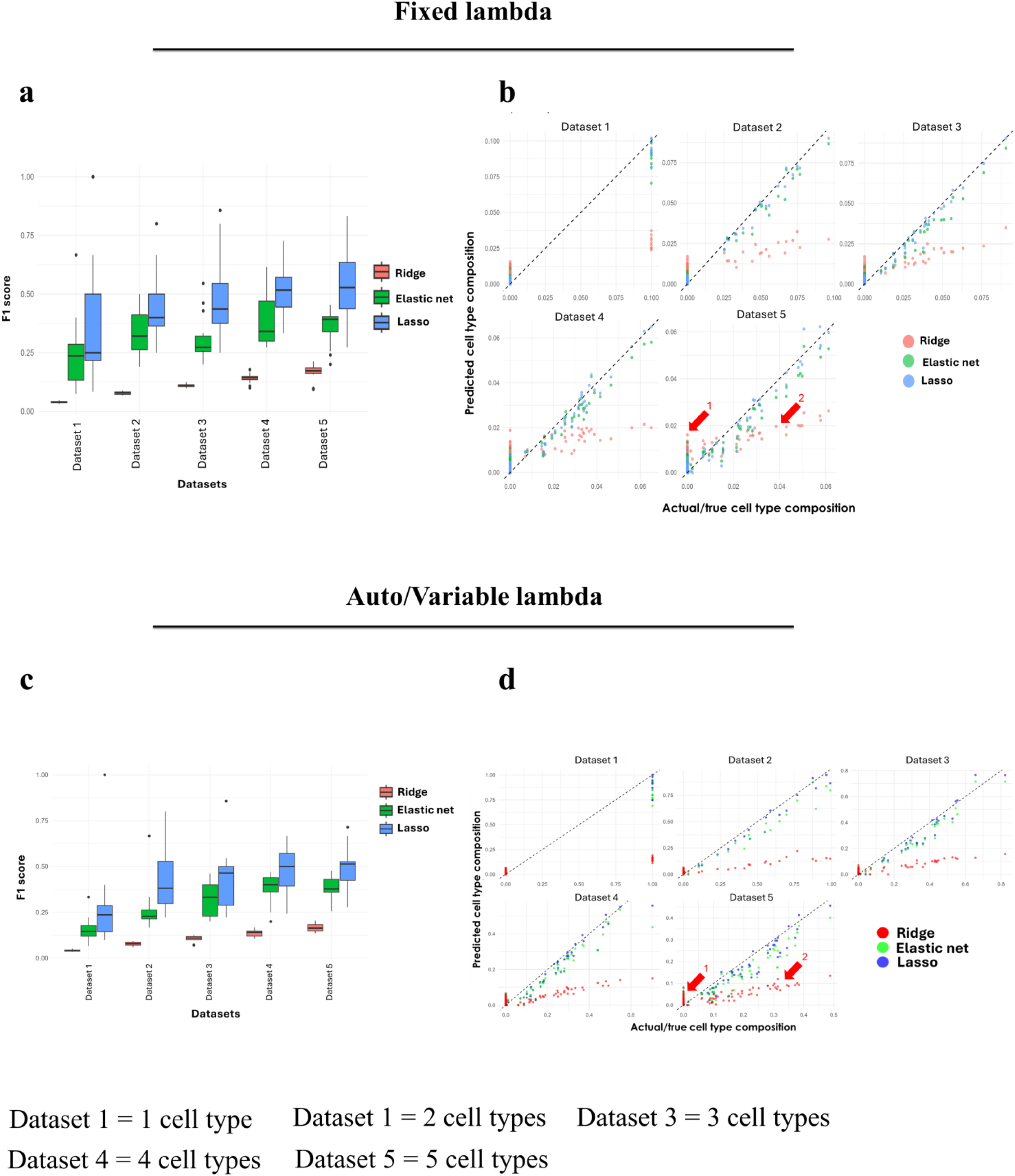
Comparing the effectiveness of shrinking techniques with fixed and automatically chosen λ on toy datasets. Fixed λ results: (a) F1 scores on toy datasets, with each dataset containing a different number of cell types using the fixed-λ approach. (b) Scatter plots of true versus predicted cell types for each dataset using Lasso, Ridge, and Elastic Net. In the scatter plot for dataset 5, Red arrow 1 shows false positive cell types, and 2 shows irrelevant cell types mostly predicted with the Ridge method, and the lowest for Lasso. Results with automatically chosen λ: (c) F1 scores on toy datasets, with each dataset containing a different number of cell types using the auto-λ approach. (d) Scatter plots of true versus predicted cell types for each dataset using Lasso, Ridge, and Elastic Net. In the scatter plot for dataset 5, Red arrow 1 shows false positive cell types, and 2 shows irrelevant cell types mostly predicted with the Ridge method. Lasso continues to be the most stable approach to predict the most accurate cell types and keeps the false positives to the lowest.

These findings collectively demonstrate that Lasso is the best technique for accurate cell type prediction, by successfully removing non-contributing cell types. Combined results for all datasets in one scatter plot for each method can be seen in Supplementary Figure S1. Lasso regression predicted the cell types closest to the true cell types while Ridge performed poorly, thus predicting many wrong cell types; however, Elastic Net, which is a combination of Lasso and Ridge regression, gave slightly better predictions than Ridge.

### Applying the approach to annotated single-cell RNA seq (scRNA) data

Having explored the accuracy of our method on toy datasets, we next applied it to a well-studied single-cell RNA-seq dataset from PBMCs with cell type annotations (36). Figure 3a shows a UMAP plot of the detected clusters and their original annotation. Three big clusters of cells were found, one of B cells, one of various T cells and NK cells, and one with myeloid cells, as well as a smaller cluster of platelets. Next, we attempted to predict these cell types using 2 approaches: 1) a default approach where we used SingleR with a general reference dataset, and 2) using oCELLoc to first select relevant cell types from the same reference, after which SingleR was used using those selected cell types. Figure 3b shows the results of approach 1 (general reference without oCELLoc). Since the reference included a wide variety of cell types, SingleR incorrectly predicted that most cells in the PBMC dataset were of an irrelevant cell type (“neuron”). This illustrates that the presence of irrelevant cell types can strongly impair the accuracy of cell type predictions. In contrast, using oCELLoc to pre-select cell types resulted in predicted cell types that are closer to the true annotations (Fig. 3c), with fewer irrelevant cell types. Moreover, remaining irrelevant cell types, such as tongue muscle cells, represent a minor percentage of the total population (Fig. 3d). In conclusion, the use of oCELLoc resulted in improved cell type predictions in this scRNA-seq dataset.

**Figure 3:**
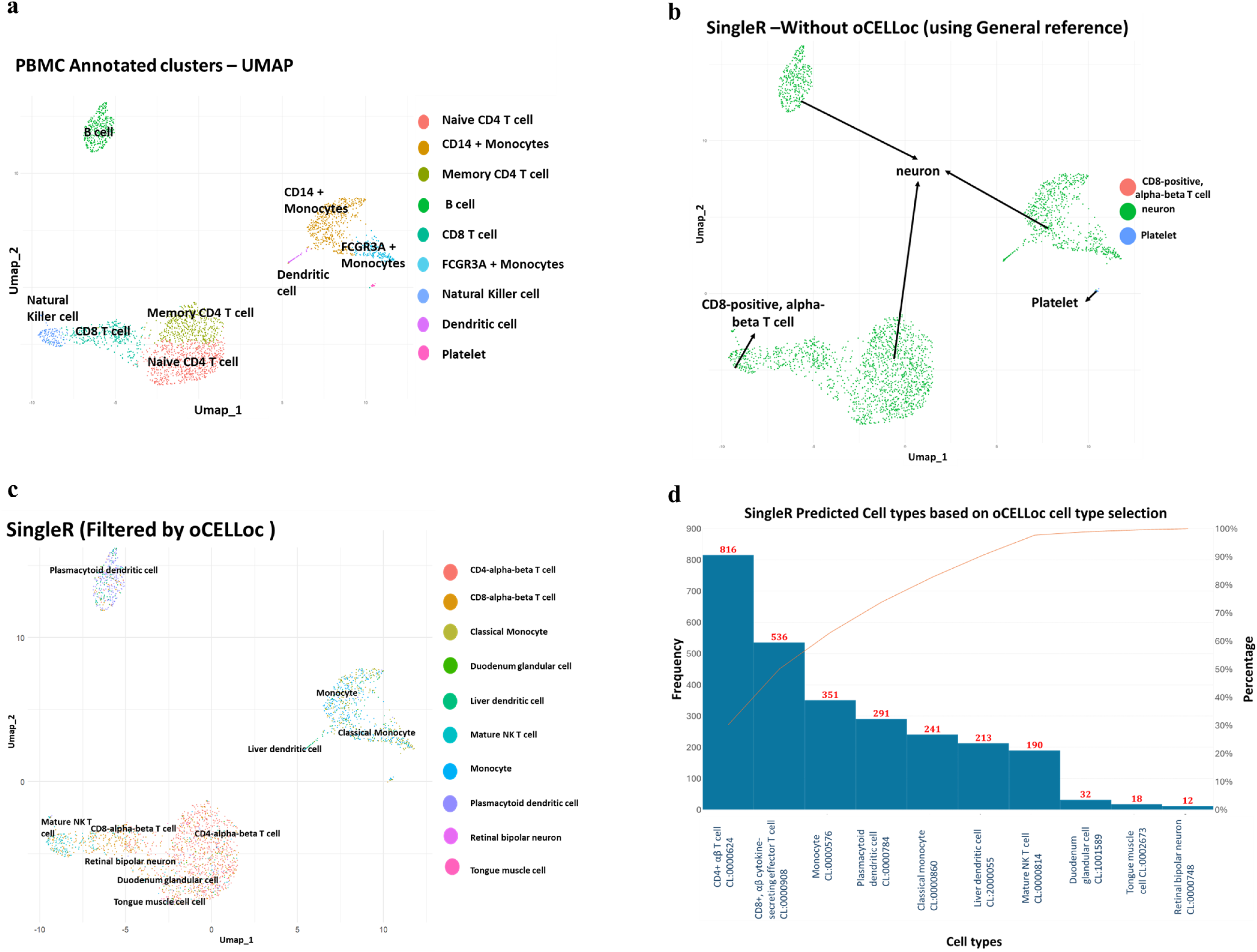
Application on scRNA datasets. (a) UMAP of the PBMC scRNA-seq dataset annotated with the actual cell types. (b) SingleR annotations utilizing a wide general reference without oCELLoc filtering. Several inaccurate labels, such as neurons and unrelated cell types, are predicted. (c) SingleR annotations following pre-filtering the reference using oCELLoc. Biologically meaningful clusters are predicted by eliminating unnecessary cell types, thus closer to actual annotations. (d) The frequency distribution of cell types that oCELLoc predicted shows that small, irrelevant cell types, such as tongue muscle cells, represent a minor percentage of the total cell types.

### Cell type selection in Visium samples using oCELLoc

We collected 2144 Visium samples, out of which 1361 were human and 783 were mouse samples. These tissues comprise a wide variety of tissues and conditions from various sources. To apply oCELLoc to such a diverse set of data samples, we prepared a large reference dataset for mouse and human, including expression patterns for tumor cells (see Methods). We applied oCELLoc on each Visium sample using Lasso regularization. In general, a spatial sample is first converted to a pseudobulk expression matrix by averaging gene expression across all spots. A broad reference matrix with pseudobulk expression of 125 cell types (170 human and 125 mouse as described above) was prepared. Each pseudobulk sample and the reference were given as input to oCELLoc, which filtered and predicted the most suitable cell types in each sample. These filtered cell types were then used as a reference to run RCTD spatial deconvolution. Resulting tissue-specific cell type prediction results for human and mouse are listed in Supplementary Table S4 and S5, respectively.

### Cell type variations in mouse and human samples

Interestingly, the model’s output was significantly impacted by both the composition and the high quality of the reference datasets. The mouse ST datasets included several healthy tissues, where patterns of cell type expression were clearer and less affected by pathological conditions, thus being more similar to the reference scRNA data. Because of this, oCELLoc consistently generated predictions for the mouse datasets that were more accurate and comprehensible, with a better degree of agreement between the anticipated and known biological composition. Figure 4a shows the top predicted cell types across mouse tissues, which closely match the expected dominant cell type. Examples for liver and heart are highlighted in Figure 4b. oCELLoc results showing the top 10 predicted cell types that include the most dominant cell types as hepatocytes in the liver and heart muscle cells in the heart, respectively Figure 4c. In contrast, the results of human samples show more complexity, likely caused by a majority of the human ST datasets originating from post-mortem, diseased, or malignant tissues, as well as the higher biological heterogeneity among patients. Predicted and expected cell types occasionally showed less harmony because of these complications, which frequently led to more varied predictions but still predicted the necessary cell types. Figure 5a shows a summary of the top predicted cell types across human tissues. Zooming in on skin and pancreas (Figure 5b), a considerable degree of variability between predicted cell types among samples can be observed. Nevertheless, in general, dominant cell types predicted in each tissue are in general biologically relevant (Figure 5c). In summary, these findings demonstrate that oCELLoc can effectively adapt to complex spatial transcriptomics data and identify subsets of contributing cell types, even in heterogeneous or noisy human tissue samples. However, for some samples, human supervision is necessary to obtain accurate cell type predictions. Although our reference datasets contain gene expression patterns of a large variety of cell types, they do not accurately represent the full variety of different states that can be occupied by different cell types, such as, for example, activated immune cells. As a result, in general, the accuracy of predictions is higher for mouse samples, which are relatively more similar to the mouse reference. In contrast, human samples are often obtained post-mortem of from patients suffering from medical conditions. It is reasonable to assume that our human reference data does not align as well with such samples.

**Figure 4:**
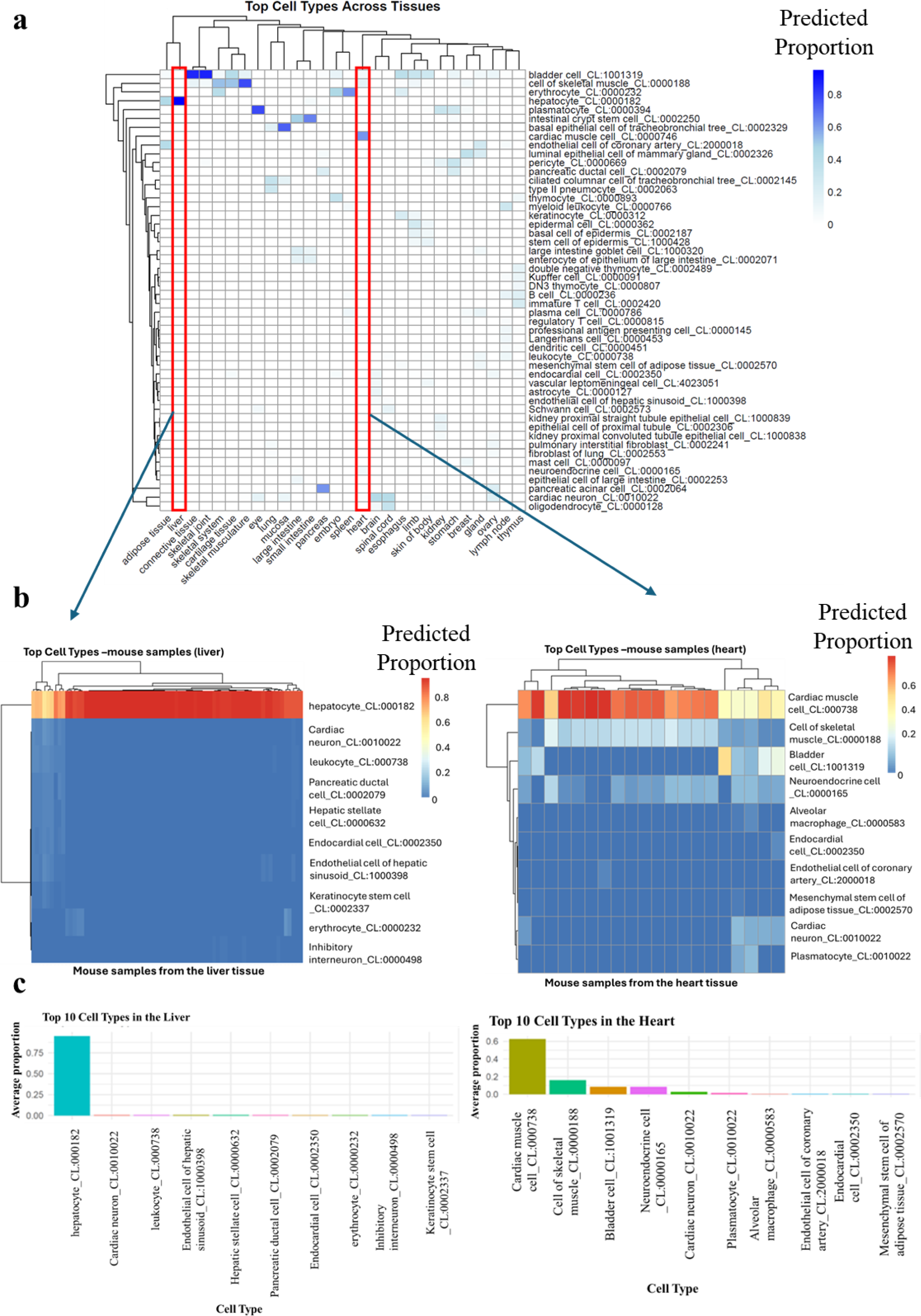
Application of oCELLoc on mouse ST datasets. (a) Heatmap showing the top contributing cell types predicted by oCELLoc across several tissues, with a scale from 0 (light grey) to 1 (dark blue), 1 showing the highest concentration of the contributing cell types and close to zero being the lowest. (b) Heatmaps of the cell types in the liver and heart tissues of mouse samples. Red color represents the highest proportion of cell type, yellow shows moderate or between 0 and 1, while blue shows lower proportions of cell types. (c) Top 10 cell types in liver and heart tissues.

**Figure 5:**
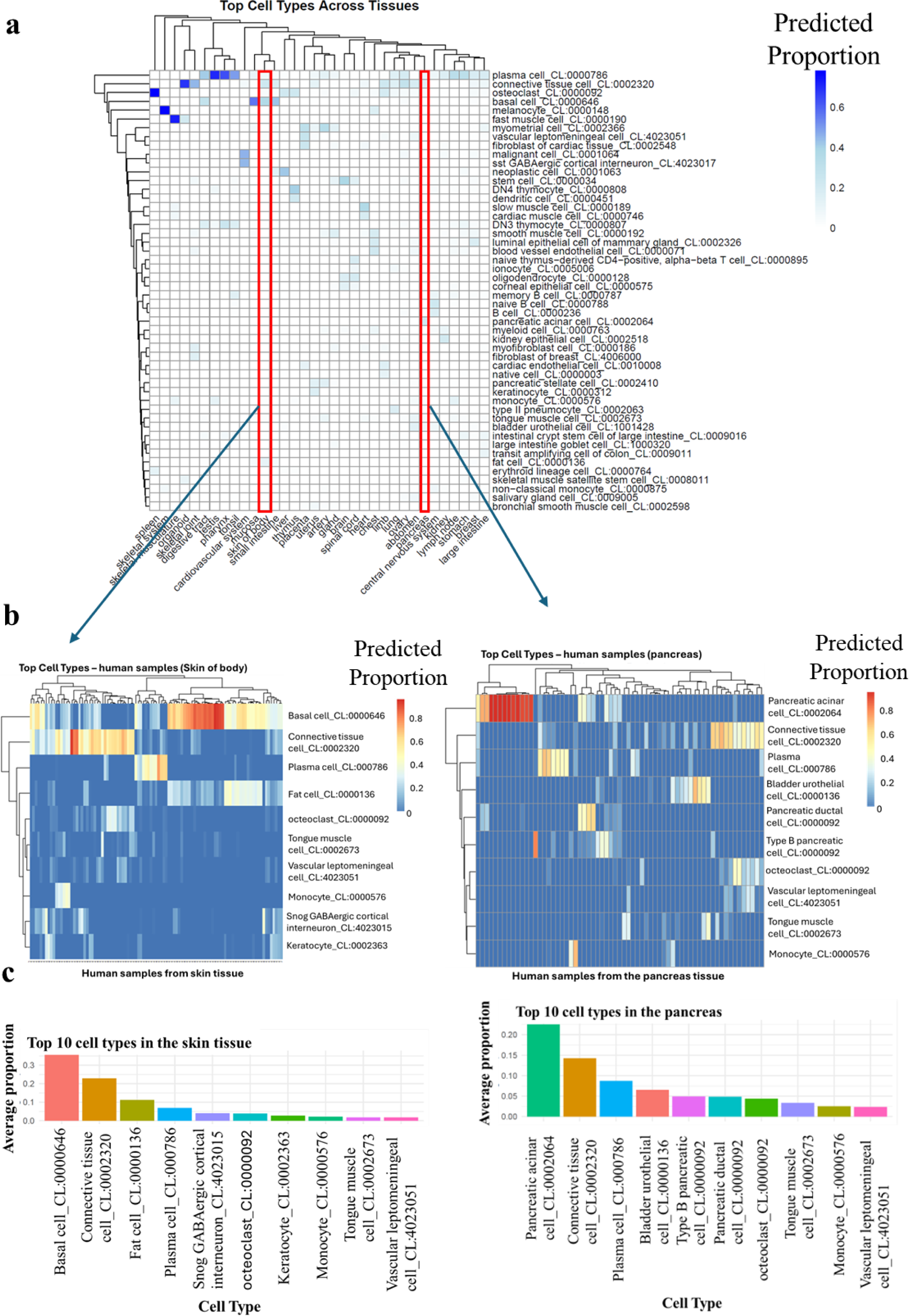
Application of oCELLoc on human ST datasets. (a) Heatmap showing the top contributing cell types predicted by oCELLoc across several tissues. With a scale from 0 (light grey) to 1 (dark blue), 1 showing the highest concentration of the contributing cell types and close to zero being the lowest (b) Heatmaps of cell types in the skin and pancreas tissues of mouse samples. Red color represents the highest proportion of cell type, yellow shows moderate or between 0 and 1, while blue shows lower proportions of cell types. (c) Top 10 cell types in human skin and pancreatic tissues.

In oCELLoc, the λ value controls the amount of shrinkage, and therefore the number of cell types retained in the result. In samples where a higher (i.e., close to 1) λ value was picked, fewer cell types had non-zero coefficients (Supplementary Figure S2). In this study, we set the minimum cell types to be included as 4 and the maximum to be 10. However, users can set these values depending on their preferences.

## Discussion

In this study, we present oCELLoc, a computational approach that identifies the most suitable cell types in scRNA or ST transcriptomic datasets and directs the creation of a minimal, high-relevance reference for downstream cell type prediction or deconvolution. We showed that oCELLoc can accurately recover known cell type compositions, lower reference complexity, and provide better downstream annotation by benchmarking on toy datasets, single-cell generated pseudobulk, and ST samples (human and mouse).

We illustrated the accuracy and remaining issues of oCELLoc through applications on toy datasets and real single-cell and ST data. oCELLoc accurately predicted cell types in toy data. In each application, oCELLoc was able to successfully identify crucial cell types. A better precision between reference and target data was seen in the predictions for mouse samples taken from healthy tissues, which were noticeably more stable and consistent. On the other hand, because of higher biological heterogeneity and technical noise, projections for human samples, which are frequently taken from diseased or post-mortem tissues, showed more fluctuation. This emphasizes how crucial model diagnostics like coefficient histograms and cross-validation curves are for directing parameter selection and guaranteeing interpretability. Together, these results demonstrate that oCELLoc is a useful pre-deconvolution approach that, by eliminating unnecessary or confusing cell types from sizable reference datasets, improves the interpretability, effectiveness, and precision of downstream cell type prediction of deconvolution.

### Comparison with existing methods and the landscape of spatial deconvolution

In contrast to other related approaches, oCELLoc aims to be a filtering/selection layer upstream of cell type prediction tools that assume a correct reference is provided as input. In this way oCELLoc assists subsequent inference on a refined panel and reduces the noise caused by irrelevant cell types. However, the quality of the reference, the match between the reference and the target (in terms of species, physiological/disease state, and batch effects), and gene selection or normalization techniques all affect how well any deconvolution technique performs (37).

### Conclusion and future prospects

Although oCELLoc has many advantages, there is always room for improvement. The quality and coverage of the dataset used for reference have a significant impact on its performance; if important cell types or states are absent or under-represented, the model may reject them or replace them with closely related types. Additionally, the technique is susceptible to λ tuning and regularization settings; however, this is lessened by cross-validation. In heterogeneous human samples, different λ values may produce different cell type subsets, and no golden standard dataset is available. Furthermore, the linear mixture model of expression used by oCELLoc might not adequately account for nonlinearities resulting from RNA diffusion or cell-cell interactions.

The framework could be improved in several ways in the future. Consistency between adjacent regions may be enhanced by incorporating nearest neighbor information from existing literature, and adaptive reference improvement may repeatedly improve the chosen reference depending on any remaining errors. These processes could be unified and accuracy increased by creating a joint selection and deconvolution model that resembles sparse Bayesian formulations (15). Furthermore, managing huge reference atlases will require high-performance and scalable computation, and additional benchmarking across other tissues and disease states, such as cancer, fibrosis, and developmental systems, will validate the generalizability of the model (38). Lastly, the integration of oCELLoc into end-to-end deconvolution workflows (like Cell2location, RCTD, or SPOTlight) will help the spatial transcriptomics community adopt and implement the approach more widely.

In conclusion, oCELLoc fills a critical but frequently overlooked need in spatial transcriptomics deconvolution: the space of potential cell types must be pruned before inference. oCELLoc consistently finds a minimal set of cell types that best explains a target dataset through the use of shrinkage (Lasso) and cross-validation. This allows for more targeted downstream deconvolution or annotation. Our tests show that this method increases flexibility across data modalities, reduces the risk of overfitting, and enhances interpretability. Even though there are still difficulties, particularly with diseased human tissues and spatial coherence modelling, we see oCELLoc as a broad tool in the spatial transcriptomics toolbox and encourage the community to expand on it in subsequent methodological developments.

## Acknowledgements

The authors would like to thank Y. Harada for secretarial assistance.

